# Alterations in gut microbiota linked to provenance, sex, and chronic wasting disease in white-tailed deer (*Odocoileus virginianus*)

**DOI:** 10.1101/2021.01.11.426270

**Authors:** David Minich, Christopher Madden, Morgan V. Evans, Gregory A. Ballash, Daniel J. Barr, Keith P. Poulsen, Patricia M. Dennis, Vanessa L. Hale

## Abstract

Chronic wasting disease (CWD) is a fatal, contagious, neurodegenerative prion disease affecting both free-ranging and captive cervid species. CWD is spread via direct or indirect contact or oral ingestion of prions. In the gastrointestinal tract, prions enter the body through microfold cells (M-cells), and the abundance of these cells can be influenced by the gut microbiota. To explore potential links between the gut microbiota and CWD, we collected fecal samples from farmed and free-ranging white-tailed deer (*Odocoileus virginianus*) around the Midwest. Farmed deer orignated from farms that were depopulated due to CWD. Free-ranging deer were sampled during annual deer harvests. All farmed deer were tested for CWD via ELISA and IHC, and we used 16S rRNA gene sequencing to characterize the gut microbiota. We report significant differences in gut microbiota by provenance (Farm 1, Farm 2, Free-ranging), sex, and CWD status. CWD-positive deer from Farm 1 and 2 had increased abundances of *Akkermansia*, *Lachnospireacea* UCG-010, and RF39 taxa. Overall, differences by provenance and sex appear to be driven by diet, while differences by CWD status may be linked to CWD pathogenesis.

## Introduction

Chronic wasting disease (CWD) is a fatal, contagious, neurodegenerative prion disease affecting both free-ranging and captive cervid species, including white-tailed deer (*Odocoileus virginianus*), mule deer (*Odocoileus hemionus*), elk (*Cervus elaphus elaphus*), and moose (*Alces alces*). First identified in Colorado, USA in the 1960s, CWD was given the designation as a transmissible spongiform encephalopathy (TSE) in 1978 ^1,2^. Other TSEs include bovine spongiform encephalopathy, transmissible mink encephalopathy, kuru, and variant and sporadic Creutzfeldt-Jakob Disease (CJD) ^1^. Since the 1960s, CWD has spread across North America and has been identified in cervids in 26 states ^3^. Outside of the United States, CWD has been documented in Korea, Canada ^1^, and Norway ^4^. Clinical signs of CWD include progressive weight loss, altered posture, head tremors, ataxia, and polydipsia and polyphagia^1^. Pathologically, CWD causes spongiform lesions within the central nervous system caused by an abnormal, diseased isoform (PrP^CWD^) of the normal cellular prion protein (PrP^C^). PrP^C^ is typically composed of multiple alpha-helices, but the abnormal isoform undergoes a transformation into a beta-sheet conformation, making it resistant to proteases, high temperatures, and standard disinfection protocols ^1^. The extreme hardiness of the diseased prion, as well as an incubation period ranging from 18 months to 5 years ^1^, makes CWD extremely challenging to control and manage.

CWD is commonly shed in the saliva, urine, feces, and skin and is spread via direct or indirect contact with infectious prions and environmental fomites ^5^. There is evidence that after oral ingestion and passage into the intestinal tract, prions enter the body through microfold cells (M-cells) ^6,7^. M-cells are specialized cells found in lymphoid follicles, the appendix, mucosal associated lymphoid tissue (MALT), and in the follicle-associated epithelium (FAE) of Peyer’s patches in the gut ^8^. M-cells are considered the gatekeeper of the intestine, as they continuously sample and internalize material from the lumen of the intestine via transcytosis to the underlying lymphoid tissue in the Peyer’s patch for initiation of mucosal and systemic immune responses ^7–9^. Studies in mice have shown that after oral entry of a TSE agent, prions initially accumulate in Peyer’s patches and mesenteric lymph nodes in the gut ^7,10^. Increased M-cell abundance has been linked to an increased susceptibility to orally acquired prion diseases, and the absence of M-cells at the time of oral exposure to infectious prions blocks neuroinvasion and disease development ^6^. Importantly, M-cell abundance can be influenced by microbes in the gut as well as by enteric inflammation, and M-cell induction and development has been linked to inflammatory cytokine stimulation and pathogen infection ^11–13^. Further, a 2009 study^14^ found that mice with intestinal inflammation as a result of increased levels of *Salmonella* had a significantly higher risk of prion disease. Therefore, increased abundance of M-cells in the gut due to a concurrent inflammation or due to increased levels of specific microbes, such as *Salmonella* ^14,15^, could potentially enhance uptake of prions from the gut lumen ^12^.

The gut microbiota serves as a defense system against pathogens and other disease-causing agents ^16^. Furthermore, the gut microbiome plays an important role in host immune development ^17^, neurogenesis ^18^, brain development ^19^, and microglia function in the central nervous system (CNS) ^20,21^. The gut microbiome has also been linked to human neurologic conditions via the “gut-brain axis ^21^.” Both Parkinson’s Disease (PD) and Alzheimer’s Disease (AD) have similarities to prion diseases and involve abnormal protein aggregates and protein misfolding occurring in the brain, including a conversion of alpha-helical structures to beta-sheet structures in PD ^22–26^. As a result of these similarities, the “prion hypothesis” suggests that PD is a prion-like disease ^27^. Studies indicate a critical relationship between the gut microbiota and neurologic diseases, including PD, AD, amyotrophic lateral sclerosis (ALS), and autism ^21,28–32^. In a 2016 study, alpha synuclein-overexpressing mice (a mouse model for PD) treated with antibiotics had an altered gut microbiota and exhibited reduced brain pathology and motor deficits, identifying direct links between alterations in the gut microbiota and brain pathology associated with PD. Further, microbial colonization of germ-free mice with stool samples from patients with PD resulted in the disease-typical protein-misfolding-mediated motor deficits ^31^. Although there is growing evidence for the role of gut microbes in neurologic diseases, there has been very little work examining the role of gut microbiota in prion diseases and no published studies, to our knowledge, on gut microbial communities and chronic wasting disease.

In this study, we used 16S rRNA gene sequencing to examine the gut microbiota of white-tailed deer (*Odocoileus virginianus*) from two deer farms (breeding facilities) that were depopulated due to the presence of CWD. Additionally, we characterized the gut microbiota of free-ranging white-tailed deer harvested from Cleveland Metroparks in northeast Ohio as part of its deer population management program. Based on previous studies that have reported differences in the gut microbiota of wild and captive ruminants, including deer ^33^, we hypothesized that microbial communities would differ between deer by provenance (Farm 1, Farm 2, and Free-ranging) with the greatest differences being observed between farmed and free-ranging deer. Based on studies that have reported alterations in gut microbiota associated with neurologic disease, we hypothesized that we would observe differences in farmed-deer gut microbial communities by CWD status (CWD-positive, CWD non-detect).

## Methods

### Fecal Sample Collection

Per United States Department of Agruculture (USDA) regulations, all deer on Farm 1 (n=101) and Farm 2 (n=30) (Wisconsin, USA) were euthanized after a deer from each farm tested positive for CWD at harvest. Farm 1 was depopulated in May 2018, and Farm 2 was depopulated in May 2019. Post-euthanasia, deer were transported to the Wisconsin Veterinary Diagnostic Laboratory (WVDL) for CWD enzyme-linked immunosorbent assay (ELISA) testing. The WVDL is a National Animal Health Laboratory Network Level 1 laboratory and is accredited by the American Association of Veterinary Laboratory Diagnosticians. Regulatory surveillance samples were shipped to the National Veterinary Services Laboratory (NVSL) for CWD Immunohistochemistry (IHC).

Fecal samples were collected digitally from the rectum of all deer and stored on dry ice until they were transferred into a −80°C freezer. Fresh gloves were donned for sampling each deer. Samples remained at −80°C until DNA extraction was performed. All deer carcasses were disposed of after sampling via an alkaline tissue digester at the WVDL. One hundred and one deer were sampled from Farm 1; thirty deer were sampled from Farm 2 (**Table 1**).

**Table 1:** Farm 1, Farm 2, and Free-Ranging Deer Demographics.

| **Farm 1** |  |  |
| --- | --- | --- |
|  | **CWD Positive** | **CWD Non-Detect** |
| Total Number of Deer | 20 | 81 |
| Age at Depopulation (mean ± SD) | 2.59 ± 1.02 | 4.37 ± 3.43 |
| Sex (n, %) |  |  |
| Female | 1 (1) | 35 (35) |
| Male | 19 (19) | 46 (45) |
| **Farm 2** |  |  |
| Total Number of Deer | 6 | 24 |
| Age at Depopulation (mean ± SD) | 2.76 ± 0.39 | 3.22 ± 2.71 |
| Sex (n, %) |  |  |
| Female | 0 (0) | 12 (40) |
| Male | 6 (20) | 12 (40) |
| **Free-Ranging Deer** | **Total** |  |
| Total Number of Deer | 100 |  |
| Age at Depopulation (mean ± SD) | 1.70 ± 1.01 | |
| Sex (n, %) |  |  |
| Female | 50 (50) |  |
| Male | 50 (50) |  |

One hundred fecal samples were also obtained from free-ranging white-tailed deer harvested in the Cleveland Metroparks (January – March, 2018; **Table 1**), as part of a deer population management program that includes regular CWD testing. Cleveland Metroparks deer herds were tested for CWD in 2008 (125 deer), 2011 (53 deer), 2012 (50 deer), 2016 (277 deer), and 2020 (135 deer), and none were found to have detectable CWD. Harvested deer were brought to a central location within four hours of death, and a fecal sample was obtained from the rectum of each deer, placed in a sterile plastic bag, and frozen at −20°C. Samples were transferred into a − 80°C freezer within 24 hours of collection where they remained until DNA extraction.

Samples from deer on Farm 1 and 2 were collected under USDA APHIS permit #136689. Post-mortem collection of feces was deemed exempt by the IACUC.

### CWD Sample Collection and Testing

The head was removed from all farmed deer greater than one year of age and the obex region of the brainstem and medial retropharyngeal lymph nodes were collected following USDA APHIS guidelines^34^. IHC and ELISA-based testing for the abnormal prion protein were performed on the dorsal motor nucleus of the vagus nerve in the obex and medial retropharyngeal lymph nodes. For IHC testing, tissues were preserved in 10% neutral buffered formalin, embedded in paraffin, sectioned at 5 μm, mounted on slides, and examined using IHC with monoclonal antibody (Mab) F99/97.6.1^35^. Animals were considered CWD-positive if any one of the tissues examined contained detectable PrP^CWD^. Animals in which tissues did not contain detectable PrP^CWD^ were considered CWD non-detect animals.

### DNA Extraction, Amplification, and Sequencing

DNA extraction on all fecal samples was performed as follows: Approximately 0.25 grams of stool was used for each extraction with QIAamp PowerFecal DNA Kits (Qiagen, Venlo, Netherlands). Following DNA isolation, DNA concentration and purity was measured using a Qubit Fluorometer 4 (Invitrogen, Carlsbad, CA, USA) and a NanoDrop 1000 Spectrophotometer (Thermo Scientific, Waltham, MA, USA), respectively. Ethanol precipitation was performed on all DNA samples from Farm 1 to improve DNA purity using a protocol^36^ from MRC Holland (Amsterdam, Netherlands). Briefly, 4 μl of sodium acetate and 132 μL of 200 proof ethanol was added to 40 μL of the DNA. This was incubated for 30 minutes at 4°C then centrifuged for 30 minutes at 4°C. After removing the supernatant, 250 μL of 70% ethanol was added to the DNA and centrifuged for 15 minutes. The supernatant was again removed and the DNA pellet was resuspended in 40 μL of the C6 elution buffer from the PowerFecal (Qiagen) DNA isolation kits. All DNA samples were submitted for library preparation and 16S rRNA gene sequencing on an Illumina MiSeq (Farm 1 and Free-ranging: The Ohio State University Molecular and Cellular Imaging Center; Farm 2 samples: Argonne National Laboratory). Earth Microbiome Project primers (515F and 806R) were used to amplify the V4 hypervariable region of the bacterial 16S rRNA gene^37^.

### Sequence Processsing and Analysis

A total of 231 samples were submitted for sequencing. Raw, paired-end reads were processed and denoised in QIIME2 v. 2020.2^38^. Taxonomy was assigned using the SILVA 132 99% amplicon sequence variants (ASVs) database from the 515F/806R classifier^39,40^, and samples were filtered at a sequencing depth of 10,000 features. This resulted in the retention of 229 samples with the loss of 2 samples – one CWD non-detect male deer from Farm 1 and one CWD non-detect female deer from the free-ranging population. After filtering, 5,803,410 reads from 229 samples were used for analysis (average of 25,342 reads per sample). Reads per sample ranged from 10,049 to 92,179 reads. Alpha (Shannon Diversity Index) and beta diversity were analyzed using QIIME 2^38^. Beta diversity indices were compared using permutational multivariate analysis of variances (PERMANOVA) between weighted and unweighted Unifrac distance matrices. P-values were corrected for multiple comparisons using the Benjamini-Hochberg FDR correction, and values less than 0.05 were considered significant. An analysis of composition of microbes (ANCOM) was used to determine differentially abundant taxa between groups after filtering out taxa that had fewer than 10 reads and taxa that occurred in fewer than two deer. We performed ANCOMs at both the L7 and amplicon sequence variant (ASV) levels. The L7 level is roughly equivalent to a species level while an ASV is roughly equivalent to a bacterial strain and may differ from another ASV by as few as one nucleotide^41^. Multiple ASVs may be classified as a single L7 level taxa. However, deeper genome sequencing is necessary for true species and strain differentiation as this is not feasible with amplicon sequencing alone. The single CWD-positive female was not included in statistical analyses comparing CWD-positive and CWD non-detect animals to reduce any confounding introduced by sex. Sequencing data is available at NCBI Bioproject PRJNA688284.

## Results

### Microbial Composition and Diversity by Provenance and Sex

When we examined the gut microbiota of all deer (n=229), we found significant differences in gut microbial composition and diversity by provenance (Farm 1, Farm 2, Free-ranging), with farmed deer having greater microbial diversity than free-ranging deer (Unweighted UniFrac PERMANOVA *p* = 0.001, Shannon Diversity Index *q* = 6.5 x 10^−11^, Weighted UniFrac PERMANOVA *p* = 0.001; **Fig. 1a, b, Fig. 1a**). Moreover, farmed deer from both farms had more similar gut microbiota to each other than to free-ranging deer (Farm 1 to Farm 2 pseudo-F = 9, q = 0.001; Farm 1 to Free-ranging pseudo-F =38, q = 0.001; Farm 2 to Free-ranging pseudo F = 18, q = 0.001; **Fig. 1c**).

**Figure 1.**
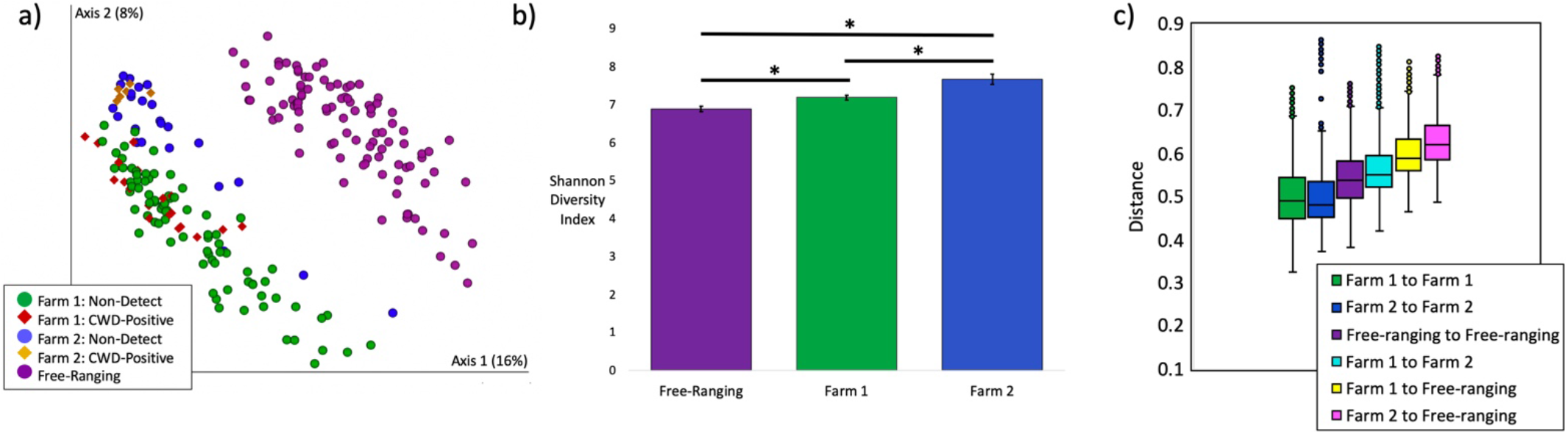
Microbial composition and diversity by provenance. a) Microbial composition (Unweighted UniFrac) differed significantly by provenance (PERMANOVA p = 0.001). Farm 1 deer are featured in green circles and red diamonds. Farm 2 deer are featured in blue circles and yellow diamonds. Free-ranging deer are featured in purple circles. b) Microbial diversity as measured by the Shannon Diversity Index differed significantly by provenance (p = 6.5 x 10^−11^). All pairwise comparisons *p < 0.001. c) Farmed deer have more similar microbial communities to each other than to free-ranging deer (Unweighted UniFrac pairwise PERMANOVA: Farm 1 to Farm 2 pseudo-F = 9, q = 0.001; Farm 1 to Free-ranging pseudo-F = 38, q = 0.001; Farm 2 to Free-ranging pseudo F = 18, q = 0.001;).

To identify microbial taxa that were differentially abundant between farmed and free-ranging deer, we combined all deer from Farm 1 and 2 – excluding CWD-positive deer – and compared these against the free-ranging deer. Through an ANCOM at the L7 (roughly species) level, we identified 82 taxa that were differentially abundant (**Supp. Table 1**). Twenty-six of these taxa were in the order Bacteroidales (phylum Bacteroidetes) and seven of these were in the family *Prevotellaceae*. The vast majority of the Bacteroidales taxas (22 of 26) were significantly increased in the farmed deer. On the other hand, free-ranging deer had significantly greater abundances of taxa (25 of 38) in the Clostridiales order (phylum Firmicutes), all of which were in the *Ruminococcaceae* and *Lachnospiraceae* families. Based on these results, we decided to compare log Firmicutes:Bacteroidetes (F:B) ratios for farmed and free-ranging deer. Log F:B ratios are associated with dietary energy harvest and higher ratios indicate greater energy extraction^42–44^. We found significantly higher F:B ratios in the free-ranging deer as compared to the farmed deer (Log F:B ratios (mean ± SE), Free-ranging: 0.39 ± 0.03; Farmed: 0.08 ± 0.02; Krustkal-Wallis *p* < 0.0001).

We also discovered significant differences in microbial composition but not diversity by sex on Farm 1 and in free-ranging deer (CWD non-detect deer only; Farm 1: Unweighted UniFrac PERMANOVA *p* = 0.008, Shannon Diversity Index *p* = 0.34, Weighted UniFrac PERMANOVA *p* = 0.003; Free-ranging: Unweighted UniFrac PERMANOVA *p* = 0.018, Shannon Diversity Index *p* = 0.53, Weighted UniFrac PERMANOVA *p* = 0.066; **Fig. 2a, b, Supp. Fig. 1b**). No significant differences in microbial composition or diversity were detected by sex on Farm 2 (CWD non-detect deer only; Farm 2: Unweighted UniFrac PERMANOVA *p* = 0.179, Shannon Diversity Index p = 0.15, Weighted UniFrac PERMANOVA *p* = 0.115; **Fig. 2a, b, Supp. Fig. 1b**. There were also no differentially abundant microbial taxa detected by sex on Farm 2. However, on Farm 1, we identified a single taxa that was significantly increased in males. This was an uncultured bacterium from the order Bacteroidales, family RF16 (ANCOM, L7 - roughly species level, *W* = 626). In the free-ranging deer population, there were multiple differentially abundant taxa by sex, with the two most differentially abundant including a microbe in the genera *Oscillibacter* and a microbe in the family *Lachnospiraceae*, genera GCA-900066575. Both of these taxa were significantly increased in males (**Supp. Table 2**).

**Figure 2.**
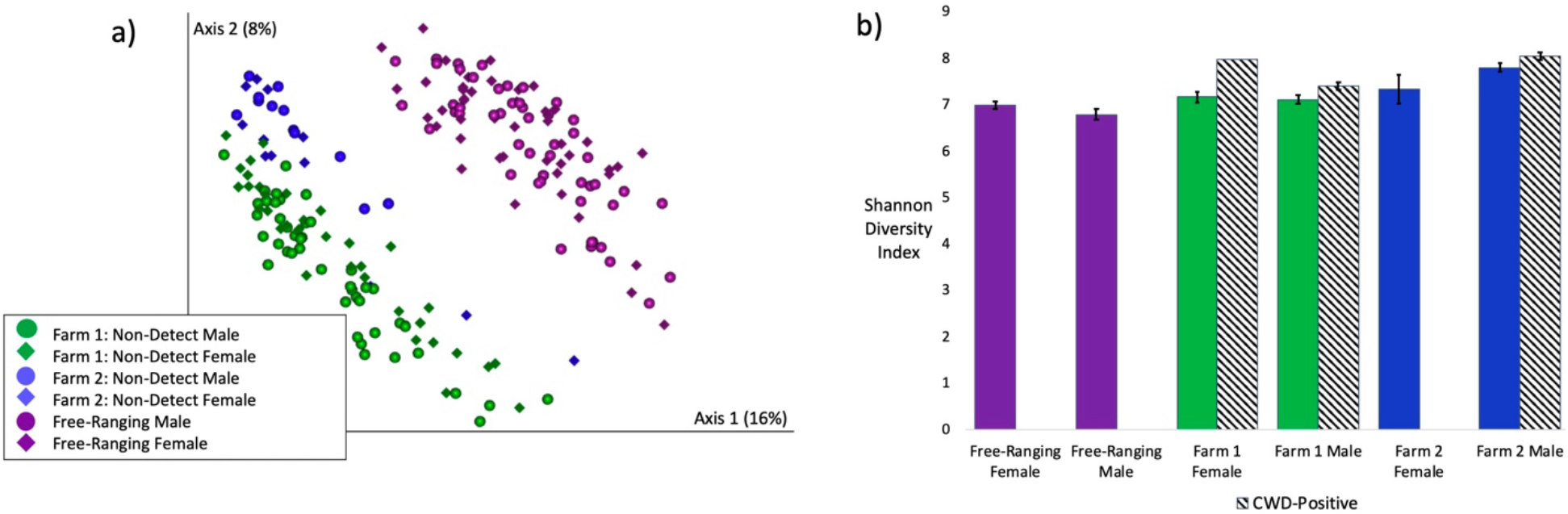
Microbial composition and diversity by sex. a) Gut microbial composition (unweighted UniFrac) differed significantly by sex on Farm 1 (PERMANOVA p = 0.008) and in Free-ranging deer (PERMANOVA p = 0.018), but not on Farm 2 (PERMANOVA p = 0.179). Farm 1 = green. Farm 2 = blue. Free-ranging = purple. Males = circles. Females = diamonds. b) Microbial diversity as measured by the Shannon Diversity Index did not differ significantly by sex (Farm 1: p = 0.34, Farm 2: p = 0.15, Free-ranging: p = 0.53) or CWD status (Farm 1: p = 0.07, Farm 2: 0.26).

### Microbial Composition and Diversity by CWD Status

Based on the microbial composition differences observed by sex and the fact that there was only one CWD-positive female in the entire data set, we opted to analyze only male deer in relation to CWD status. The single CWD-positive female deer was still included in data visualizations. Microbial composition differed significantly in CWD-positive deer on both farms (Males only; Farm 1: Unweighted UniFrac PERMANOVA p = 0.003 Weighted UniFrac PERMANOVA p = 0.011; Farm 2: Unweighted UniFrac PERMANOVA p = 0.003, Weighted UniFrac PERMANOVA p = 0.002; **Fig. 1a, Supp. Fig. 1a**). Increased microbial diversity (Shannon Index), although not significant, was also observed in CWD-positive males on both farms (Farm 1 p = 0.07; Farm 2 p = 0.26; **Fig. 2b**).

We further discovered several differentially abundant microbes at the L7 and ASV levels between CWD-positive and CWD non-detect males on both farms. (Note, multiple ASVs may be classified as a single L7, roughly species level, taxa.) On Farm 1, at the L7 level, multiple taxa were differentially abundant between CWD-positive and CWD non-detect males, the top four of which included: an uncultured bacterium from the class Bacilli (formerly Mollicutes), order RF39, increased in CWD-positive males (ANCOM *W* = 80; **Fig. 3a**); an uncultured *Paludibacter* species increased in non-detect males (ANCOM *W* = 54); an uncultured bacterium in the order Gastranaerophilales also increased in non-detect males (ANCOM *W* = 34); and a microbe in the family *Lachnospiraceae* UCG-10 increased in CWD-positive males (ANCOM *W* = 28; **Fig. 3c**) (**Supp. Table 3**). (ANCOM *W* values represent the number of times the null hypothesis is rejected in pairwise comparisons of microbial species ratios between groups. In other words, for the Bacilli RF39 L7 level taxa, the null hypothesis was rejected 80 times when comparing microbial species ratios between CWD positive and non-detect animals.)

**Figure 3.**
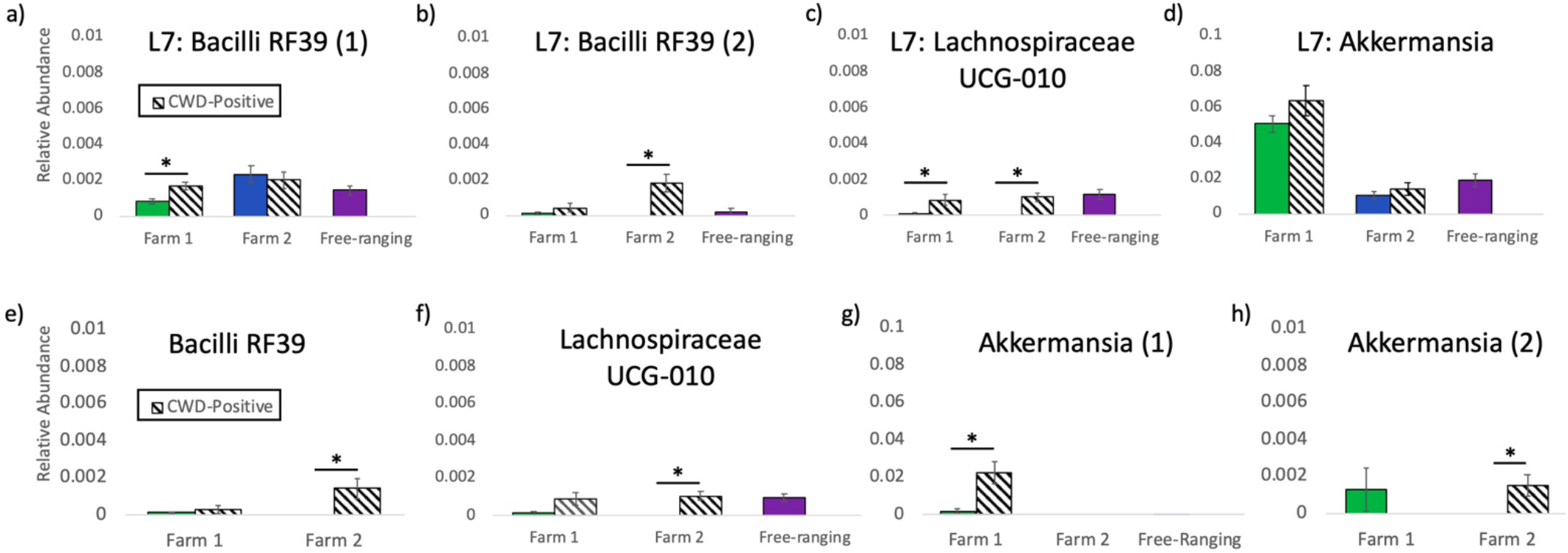
Differentially abundant microbial taxa by CWD Status. Differentially abundant taxa by CWD Status, including a) an L7 (roughly species) level taxa in the Bacilli class, order RF39, b) a second L7 level taxa in the Bacilli class, order RF39, c) an L7 level taxa in the Lachnospiraceae UCG-10 family, d) an L7 level taxa in the Akkermansia family, e) an ASV (roughly strain level) in the Bacilli class, order RF39 (formerly Mollicutes RF39), f) an ASV in the Lachnospiraceae UCG-10 family, g) an ASV in the Akkermansia family, h) a second ASV also in the Akkermansia family. Free-ranging deer (all male) did not contain any reads of the Bacilli RF39 or Akkermansia (2) ASVs.

On Farm 2, at the L7 level, two microbes were found to be differentially abundant. Both were increased in CWD-positive males and included a unidentified rumen bacterium from the class Bacilli, order RF39 (ANCOM *W* = 132; **Fig. 3b**) and a microbe from the family *Lachnospiraceae* UCG-10 (ANCOM *W* = 115; **Fig. 3c**). On Farm 2, multiple ASVs were also differentially abundant (**Supp. Table 4**), the top three of which, all increased in CWD-positive males, were an ASV from the class Bacilli (formerly Mollicutes), order RF39 (ANCOM *W* = 132; **Fig. 3e**), an ASV from the family *Lachnospiraceae* UCG-10 (ANCOM *W* = 176; **Fig. 3f**), and an uncultured ASV from the family *Akkermansia* (ANCOM *W* = 90; **Fig. 3g**). On Farm 1, at the ASV level, only one microbe was found to be differentially abundant: an ASV in the *Akkermansia* family which was increased in CWD-positive males (ANCOM *W* = 1958; **Fig. 3h**). *Akkermansia* taxa at the L7 level were not differentially abundant on either farm (**Fig. 3d**).

## Discussion

In this study, we used 16S rRNA gene sequencing to compare the gut microbiota of farmed and free-ranging white-tailed deer (*Odocoileus virginianus*). We hypothesized that deer gut microbiota would differ by provenance (Farm 1, Farm 2, and Free-ranging) and disease status (CWD-positive, CWD non-detect). Indeed, microbial composition and diversity did vary with provenance. Moreover, composition but not diversity varied with sex (Farm 1 and Free-ranging only) and with CWD status (Farm 1 and Farm 2).

### Drivers of microbial community composition by provenance

Multiple factors could contribute to the gut microbial differences we observed based on provenance, including diet, spatial proximity, host genetics, and biogeography. Diet is one of the main factors that influences gut microbial composition and diversity^45,46^. The free-ranging deer in this study had diets that primarily consisted of browse, small plants, shrubs, grasses and occassional agricultural, landscaping, and garden plants^47^. The farmed deer had access to pastures and were also fed a variety of commercial deer feeds, grains, hay, and supplemental items, including peanuts, roasted soybeans, and dandelions. As diets differed between farmed and free-ranging deer, it was not surprising that farmed and free-ranging deer had significantly different microbial communities or that there were significant differences in gut microbiota between the two farms with different feeding regimens. Multiple previous studies have also reported gut microbial differences between wild and captive animals^48–50^, including ruminants^33,51^.

Spatial proximity (or social interaction) has also been shown to influence the gut microbiota in other species: individuals with more contact share more similar gut microbiota^52^. Farmed deer sharing the same pen are likely to have increased direct and indirect contact with each other while free-ranging deer within the same herd (typically matrilineal family groups or bachelor herds) will also have more contact with each other than with non-herdmates. Host genetics can also play a role in shaping the gut microbiome^53^; although, these effects are subtle compared to other environmental factors^54^, and one previous study on white-tailed deer, albeit small (n=66), did not find any significant effects of host genetic relatedness on the gut microbiota^46^. Both farms in this study were breeding facilities and maintained a number of genetically related animals. During breeding season, a single male was commonly penned with 10-12 females for breeding. Breeders that produced high quality traits might be maintained at the farm for multiple seasons, generating several years of related offspring. Biogeography – including factors like habitat or soil type and water source – could also uniquely influence the gut microbiota of deer at each location^55^.

We hypothesized that deer gut microbial communities would differ by provenance. Specifically, we predicted that based on differing diets, host genetics, and biogeography, we would observe distinct microbial signatures in deer from each location (Farm 1, Farm 2, Free-ranging) (**Fig. 1a**). We further predicted that within locations, farmed deer would have more similar (less distant) microbiota due to more regulated diets and more limited “home ranges” as compared to free-ranging deer (**Fig 1c,** Farm 1 to Farm 1, Farm 2 to Farm 2, Free-ranging to Free-ranging). Finally, we hypothesized that the greatest differences in microbial communities would be observed between farmed and free-ranging deer since farmed deer generally share more similar diets (formulated commercial feeds, grains, hay, pasture) than free-ranging deer (**Fig. 1c**, Farm 1 to Farm 2, Farm 1 to Free-ranging, Farm 2 to Free-ranging). Our results supported each of these predictions. Although we cannot parse the individual effects of diet, spatial proximity, host genetics, and biogeography in this data set, the differentially abundant taxa identified between farmed and free-ranging deer strongly support a role for diet as a key driver of the microbial community differences we observed. Free-ranging deer consume a plant and fiber-rich diet full of shrubs and browse, while farmed deer consume a starchier diet of grains and commercial feed in addition to pasture and hay. Microbial taxa in the *Lachnospiraceae* and *Ruminococcaceae* families were increased in abundance in the free-ranging deer, while Bacteroidales taxa, like *Prevotellaceae*, were increased in the farmed deer (**Supp. Table 1**). *Lachnospiraceae* and *Ruminococcaceae* taxa are associated with plant-rich diets, and these taxa metabolize plant materials such as cellulose and hemiceullulose^50,56,57^. Bacteroidales and *Prevotellaceae* are more commonly associated with starch consumption, and in ruminants, Bacteroidales, including *Prevotella*, increase in animals on concentrate / grain diets^50,58–60^.

Firmicutes:Bacteroidetes ratios also indicated diet as a driver of differing microbial compositions between farmed and free-ranging deer. Free-ranging deer exhibited higher F:B ratios, which are associated with increased energy extraction^42^ and ferementation efficiency. In humans, increased F:B ratios are associated with obesity; in farmed ruminants, increased F:B ratios are positively correlated with average daily gain ^42,43^. In foregut-fermenting primates (which have ruminant-like digestion), wild primates exhibited higher F:B ratios than captive primates ^44^. This was attributed to the need for the wild primates to maximize energy extraction from “low-quality” food items such as fibrous plants, bark, and seeds, while captive primates, with “high quality” diets rich in soluble carbohydrates, were less dependent on efficient energy harvest^44^. Similarly, free-ranging deer gut microbiota may maximize energy extraction from a fibrous browse diet, while the grain-rich diets of farmed deer reduce the need for fermentation efficiency and create a niche for microbial taxa capable of metabolizing soluble starches and sugars. Taken together, our results suggest that diet is playing a key role in the microbial differences we observe by provenance.

### Microbial community structure by sex

Interestingly, we also identified microbial composition differences by sex on Farm 1 and in free-ranging deer. This analysis only included CWD non-detect farmed deer. No differences in microbial composition by sex were observed on Farm 2; however, Farm 2 had the smallest sample size (n=18 males, 12 females) which may have limited our power to detect these differences. Microbial community structure alterations associated with sex could be attributed to a number of factors, including differential feeding by sex or hormonal influences on the gut microbiome. We received anecdotal reports of differential feeding by sex on Farm 1 based on deer breeding and growth requirements. While we did not characterize the diet of free-ranging deer by sex in this study, a previous study on wild sheep reported differential feeding between males and females, leading to differences in gut microbiota composition between the sexes ^61^. A separate study on white-tailed deer reported that, in winter, female deer in the Midwest consumed more grass (higher quality feed) and less browse than male deer ^62^. Our samples were also collected from free-ranging deer in the Midwest during winter; thus, differential feeding could contribute to the microbial differences we observed between sexes. Male and female deer also maintain different home ranges ^63^, which can differ in vegetation – further driving potential dietary differences by sex.

Besides diet, breeding hormones have been linked to gut microbial changes in wild animals, including ground squirrels (*Spermophilus dauricus*)^64^ and black rhino (*Diceros bicornis*)^65^. It is thus possible that hormones are influencing gut microbiota in male and female white-tailed deer. Free-ranging deer were sampled January through March which corresponds to estrous cycling or pregnancy in females and post-rut (declining testosterone levels) in males^66^. Notably, our results contrast with a 2017 study on white-tailed deer that observed no differences in microbial composition between sexes; although, sampling season differed between our studies, as the 2017 study sampled deer in March and June ^46^.

Differentially abundant taxa between male and female deer included a microbe in the order Bacteroidales, family RF16 - increased in males on Farm 1; and microbes in the genera GCA-900066575 (family *Lachnospiraceae*) and *Oscillibacter* - both increased in free-ranging males. Bacteroidales and *Lachnospiraceae* taxa, discussed above, have ties to diet and energy extraction. *Oscillibacter* species increase in humans on diets high in resistant starch and low in carbohydrates ^67^, which is consistent with the browse-rich winter diet of free-ranging male deer ^62^. These differentially abundant taxa underscore the role of diet in microbial community differences observed by sex.

### Chronic wasting disease and the gut microbiota

On both farms, we observed significant differences in microbial composition in CWD-positive deer as compared to non-detect deer. Twenty-five of the 26 total CWD-positive deer across both farms were male. Previous studies in wild deer have reported that CWD prevalance is two times higher in males, and that males have a threefold greater risk of CWD infection as compared to females ^68^. These differences in infection risk and prevalence by sex are thought to be linked to increased CWD transmission amongst male social groups outside of breeding season ^68^.

Alternately, models of CWD outbreaks in captive deer predict that density-dependence and indirect transmission ^69^ play an important role in CWD spread. On at least one of the farms in this study (Farm 1), male deer were penned with females during rut (fall) and then separated into bachelor herds for the remainder of the year. As such, both transmission through male social groups and indirect, density-dependent transmission (in bachelor pens) could have played a role in the predominantly male infections observed in farmed deer. Because of this skew by sex, we opted to analyze only males in relation to CWD status. This within-farm, male-only analysis mitigated potential gut microbial confounders, including sex, diet, and biogeography.

Differentially abundant microbial taxa common across both farms and increased in CWD-positive animals included: two different microbes in the class Bacilli, order RF39 (formerly Mollicutes RF39) – one increased on Farm 1 and one increased on Farm 2; a microbe in the family *Lachnospiraceae* UCG-10; and two different ASVs in the *Akkermansia* family – one increased on Farm 1, and one increased on Farm 2 (**Fig. 3**). The fact that these three taxa (RF39, *Lachnospiraceae* UCG-10, *Akkermansia*) emerged as CWD-associated on two independently run farms over 100 miles apart is intriguing and merits further attention. In a previous study, RF39 was found to be increased in a mouse model of the relapse-remitting form of multiple sclerosis (MS) ^70^, which is a disease that shares many features with prion diseases, including CJD ^71^. Taxa in the Bacilli (formerly Mollicutes) class have been associated with CWD in other studies ^72^. Specifically, Bastian et al. reported the presence of *Spiroplasma* DNA in the brains of eight out of ten sheep with scrapie, six out of seven cervids with CWD, and two humans with CJD ^72^. All matched normal sheep, cervid, and human brains were negative for *Spiroplasma* DNA. However, no *Spiroplasma* could be detected in a hamster model of scrapie ^73^. Further, Bastian et al. induced spongiform changes in the brains of deer, sheep, and goats inoculated intracranially with *Spiroplasma* species ^74^. This could not be replicated in a subsequent study on neonatal goats ^75^; although, differences in methodology between the studies was noted. *Spiroplasma* species are not in the order RF39.

Besides RF39, we also observed an increase in *Lachnospiraceae* UCG-010 in CWD-positive animals on both farms at the L7 level (**Fig. 3b, c**). *Lachnospiraceae* taxa have been reported in other studies on wild and captive deer gut microbiota ^76,77^. *Lachnospiraceae* has also been noted in association with neurologic diseases. However, it is decreased, rather than increased, in several studies on Parkinson’s disease (PD), and this decreased abundance is associated with more severe cognitive and motor impairments ^78,79^. Decreases in *Lachnospiraceae* have also been observed in Alzheimer’s Disease (AD) and amyotrophic lateral sclerosis (ALS) ^28,80^. Moreover, multiple studies highlight the ability of Lachnospiraceae species to produce butyrate which helps maintain the epithelical barrier ^81,82^. However, *Lachnospiraceae* family taxa have also been associated with type 2 diabetes ^82^ and intestinal inflammation^83^.

Like *Lachnospiraceae*, *Akkermansia* taxa are commonly associated with health^84^ and even touted as promising probiotics ^84,85^; although, more recent evidence has promoted caution in defining *Akkermansia* as exclusively a “good bug” ^86^. In fact, the mucin-degrading *Akkermansia* is reportedly increased in multiple neurologic diseases, including PD, multiple sclerosis, and AD ^32,87–90^, although it has also been shown to reduce pathological alterations (amyloid beta-protein accumulation) and cognitive impairments in one mouse model of AD ^91^. *Akkermansia* has additionally been associated with fasting or malnutrition, as it can utilize host mucin as its sole energy source while other microbes require dietary substrates consumed by the host ^50,92^. A single *Akkermansia* ASV was significantly increased on Farm 1, while a different *Akkemansia* ASV was increased on Farm 2 (**Fig. 3a,b,g,h**), suggesting potential species or strain differences in these taxa by Farm. Future work with deeper sequencing is necessary to assess true species and strain level differences between farms.

We hypothesized that we would observe differences in gut microbial communities by CWD status, and our results support this hypothesis. However, how and why these three taxa (RF39, *Lachnospiracea* UCG-010, *Akkermansia*) are associated with CWD are the next important questions to answer. Do these taxa contribute to a gut environment that is more permissive to orally-ingested prions? *Akkermansia*, for example, can degrade mucin, thinning the protective mucus barrier that lines the gut. In concert with a pro-inflammatory *Lachnispiraceae* species, these microbes could create an inflammatory environment that induces colonic M-cells ^11,14,15,93^, enhancing susceptibility to prion disease ^6,94^. Gut inflammation has also been linked to the progression of neurodegenerative disease including AD and PD ^30–32^. Alternately, are these taxa increased as a result of prion disease? Early clinical signs of CWD can include behavioral and locomotive changes followed by eventual wasting and weight loss^1^. Subtle behavioral changes could conceivably alter diet and drive dietary differences in the gut microbiota between deer with and without CWD. *Akkermansia* can also thrive in the face of malnutrition as it only needs host mucin to survive; therefore, *Akkermansia* could increase in a host that is consuming less food. Finally, could these taxa be providing protective effects in the presence of a prion disease? This seems less likely from an evolutionary perspective, but *Lachnospireacea* and *Akkermansia* are associated with many health benefits, and increased relative abundances of these species are associated with protection against metabolic diseases and reduced pathological changes in AD ^28,82,86,91,95^. Bacilli (e.g. RF39 - formerly in phylum Tenericutes, now in Firmicutes) have also been posited to play a protective role in the gut as they are decreased in relative abundance in the presence of DSS-colitis ^96^. It is important to note that all three taxa were also observed in the free-ranging deer at varying and often comparable levels to the levels observed in CWD-positive farmed deer (**Fig. 3**). Given the significant differences in microbial composition and diversity between farmed and free-ranging deer, these results are more challenging to interpret but suggest that, while the differentially abundant taxa in CWD-postive animals may play a role in CWD pathogenesis, these results need to be interpreted carefully and within context.

This study represents the first investigation, to our knowledge, of white-tailed deer gut microbiota in relation to CWD. We acknowledge several limitations to the present study. First, while our results suggest that differential diets are the major driver of microbial community differences by provenance and sex, we cannot explicitly rule out the potential effects of spatial proximity, host genetics, or biogeography. Second, as farmed and free-ranging deer had significantly different microbial communities, we cannot be certain that microbial composition differences observed in farmed deer based on CWD-status are generalizable to free-ranging deer. Third, the free-ranging deer in this study were not explicitly tested for CWD but were presumed CWD non-detect based on extensive CWD testing on Cleveland Metroparks deer herds in years antecedent and subsequent to 2018. Further, until December 2020, CWD had never been detected in any free-ranging deer in the state of Ohio. Fourth, microbial composition is not representative of microbial function ^97^, and future studies using shotgun metagenomics and metabolomics are warranted to capture function. Fifth, while *Akkermansia*, RF39, and *Lachnospiraceae* UCG-010 are associated with CWD, further work is needed to clarify if these differences preceded or succeeded disease. Finally, fecal samples from Farm 1 and free-ranging deer underwent library preparation and sequencing on an Illumina MiSeq at The Ohio State University Molecular and Cellular Imaging Center, while fecal samples from Farm 2 underwent library preparation and sequencing on an Illumina MiSeq at Argonne National Laboratory. While differences between laboratories and sequencing facilities can lead to differing results in microbiome studies ^98^, we limited these effects by using the same methodology and kits (Qiagen PowerFecal) for all DNA extractions, the same region and primers for sequencing (V4 -515F and 806R), and all sequencing data was combined and underwent sequence processing and taxonomy assignment together. Further, our results by sex and CWD status would not be affected, as these results were analyzed independently for each location (Farm 1, Farm 2, Free-ranging).

In conclusion, we report differences in gut microbiota in white-tailed deer by provenance (Farm 1, Farm 2, Free-ranging), sex, and CWD status. Differences by provenance and sex are likely driven by diet, while differences by CWD status are more challenging to interpret and include increased abundances of *Akkermansia*, *Lachnospireacea* UCG-010, and RF39 taxa in CWD-positive deer. Priorities for future research include determing how these taxa play a role in CWD susceptibility or pathogenesis, characterizing the gut microbiota of free-ranging cervids with CWD, and assessing M-cell presence and abundance in CWD-positive and CWD non-detect animals to elucidate potential relationships between gut microbiota, M-cells, and chronic wasting disease.

## Author Contributions

DM collected the samples from the farmed deer, performed DNA extraction on all samples, assisted with data analysis of sequencing results using QIIME 2, and drafted the manuscript.

CM assisted with DNA extraction of all samples, DNA purification of Farm 1 samples, coordinated laboratory activities, and assisted with editing of the manuscript.

ME assisted with data analysis and processing of sequencing results using QIIME 2 and provided feedback on the manuscript.

GB provided expertise on neuropathology in cervids and feedback on the manuscript.

DB and KP facilitated CWD testing and sample collection from farmed deer, provided expertise on CWD in cervids, and provided feedback on the manuscript.

PD facilitated sample collection from free-ranging deer and provided feedback on the manuscript.

VLH conceived the presented idea and designed and directed the study. Additionally, VLH assisted with analysis of sequencing results, manuscript preparation, and figure generation.

## Acknowledgements

We acknowledge the many individuals at WVDL, USDA, Cleveland Metroparks, and the veterinary students at the University of Wisconsin for assistance with sample collection from farmed and free-ranging deer. We further acknowledge the Ohio State University Infectious Diseases Institute, Department of Veterinary Preventive Medicine, and the Cliff M. Monahan Summer Research Fellowship for their support of this project.

## Conflicts of Interest

No authors report any conflicts of interest.

---

**Supplemental Figure 1.**
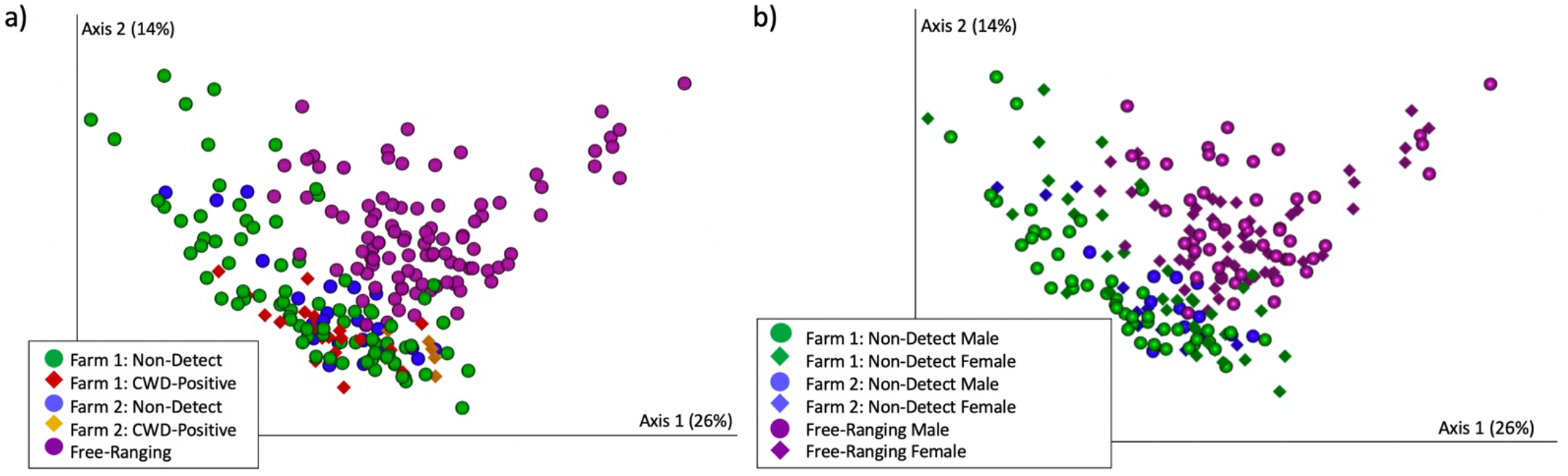
Microbial composition by provenance and sex. a) Microbial composition (weighted UniFrac) differed significantly by provenance (PERMANOVA p = 0.001). Farm 1 deer are featured in green circles and red diamonds. Farm 2 deer are featured in blue circles and yellow diamonds. Free-ranging deer are featured in purple circles. b) Gut microbial composition (weighted UniFrac) differed significantly by sex on Farm 1 (PERMANOVA p = 0.003), trended toward significance in free-ranging deer (PERMANOVA p = 0.066), but did not differ significantly on Farm 2 (PERMANOVA p = 0.115). Farm 1 = green. Farm 2 = blue. Free-ranging = purple. Males = circles. Females = diamonds

**Supplemental Table 1:**
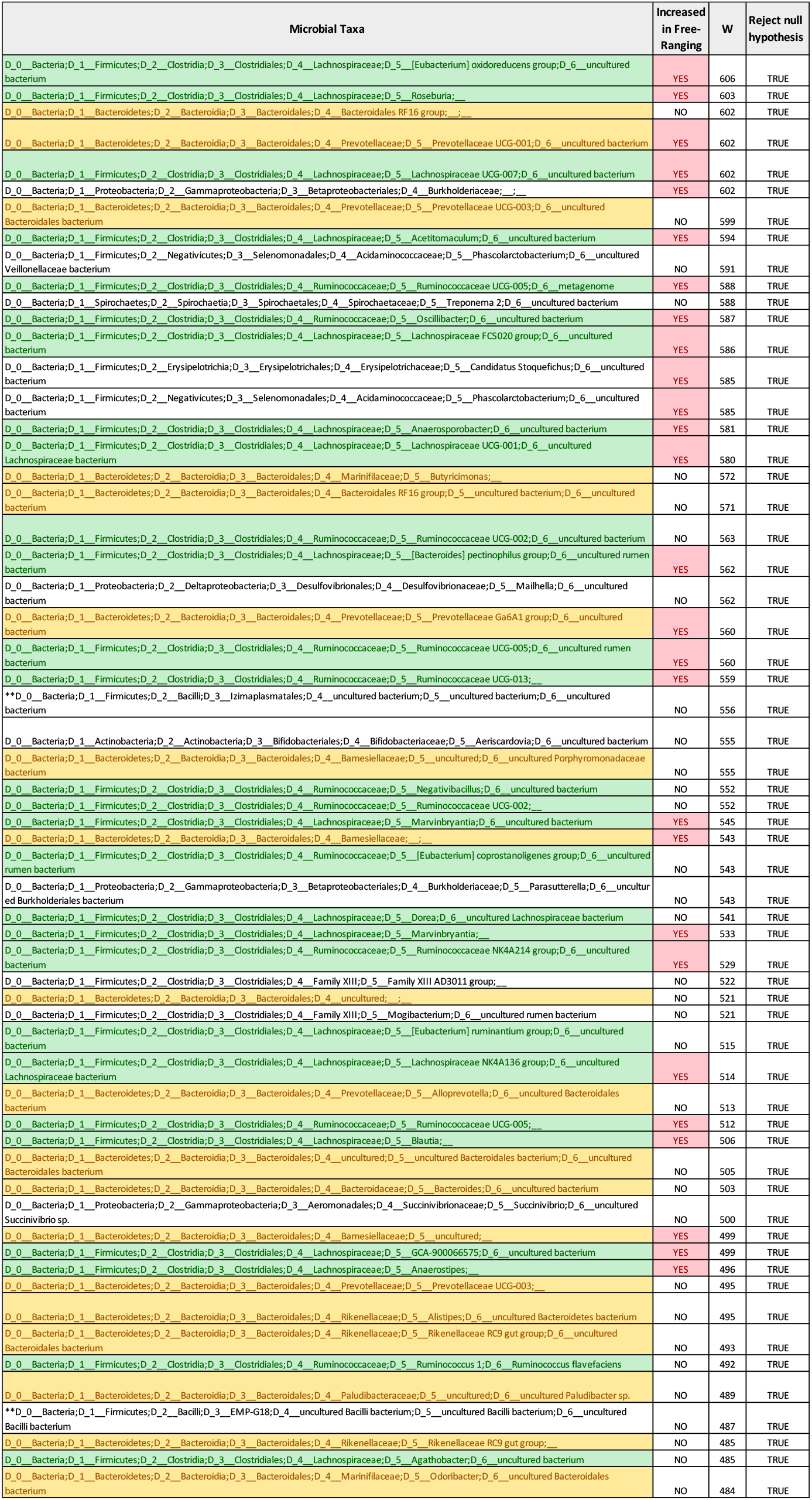
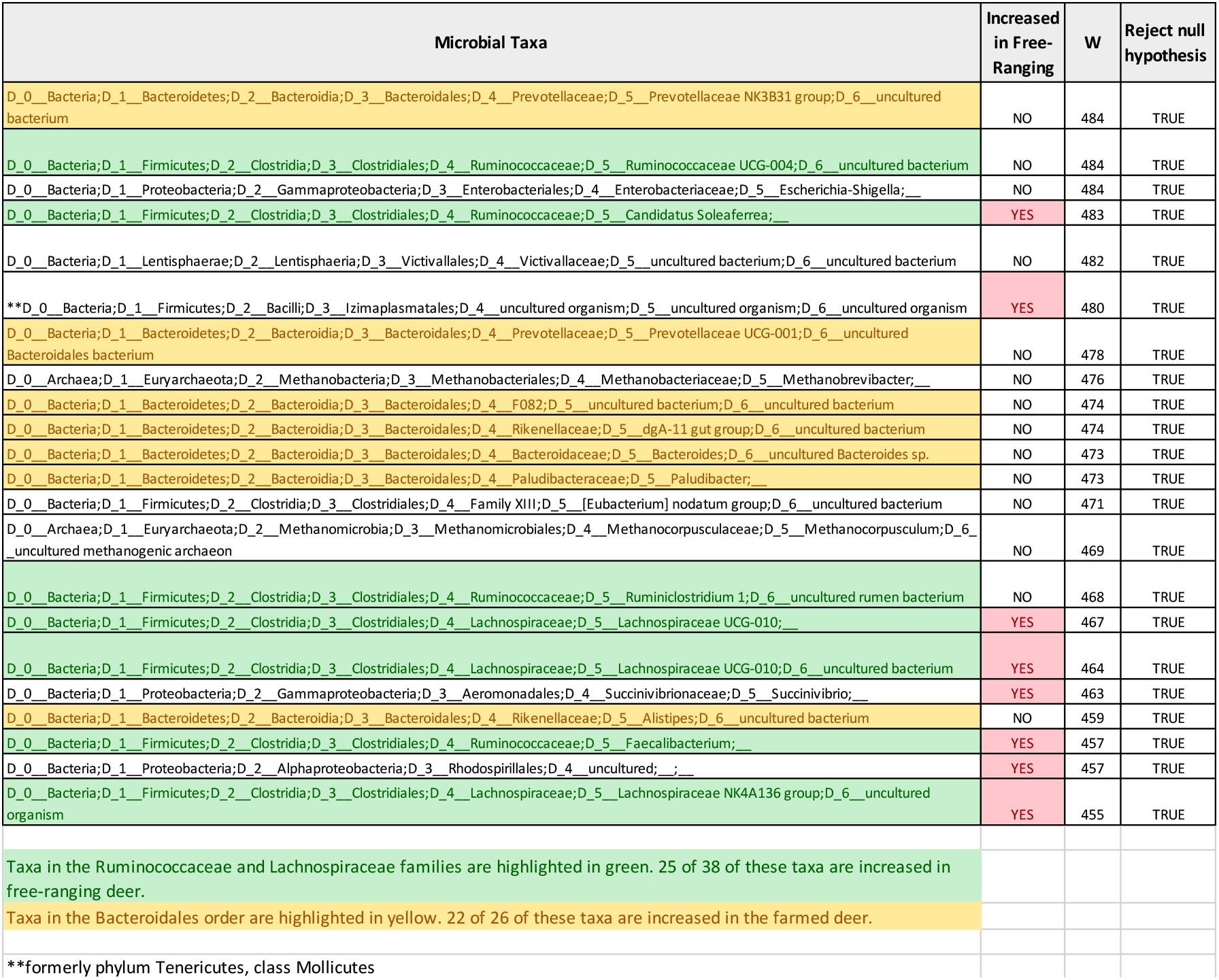
Differentially abundant taxa (ANCOM) between farmed and free-ranging white-tailed deer.

**Supplemental Table 2:**
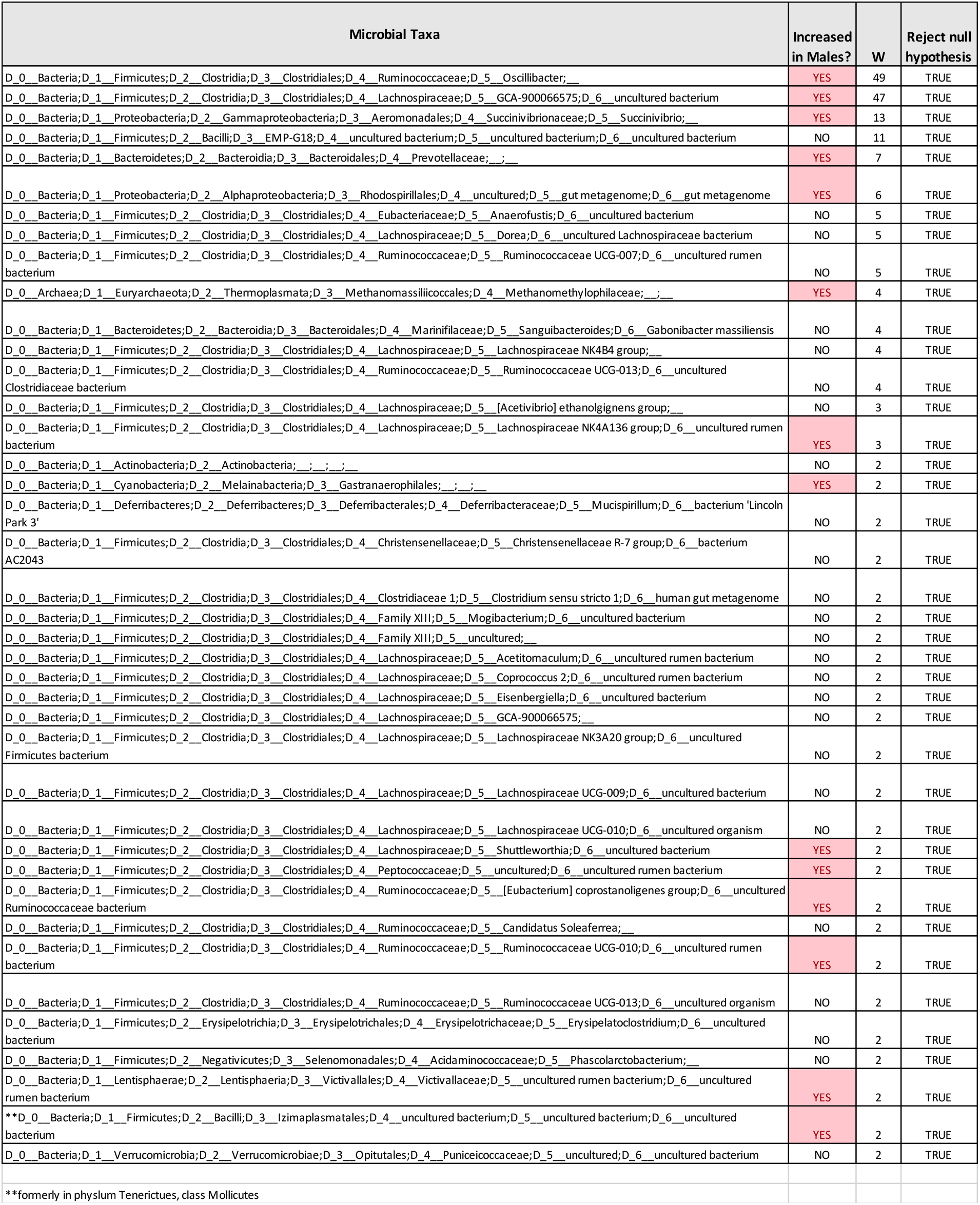
Differentially abundant taxa (ANCOM) by sex in free-ranging white-tailed deer.

**Supplemental Table 3:**
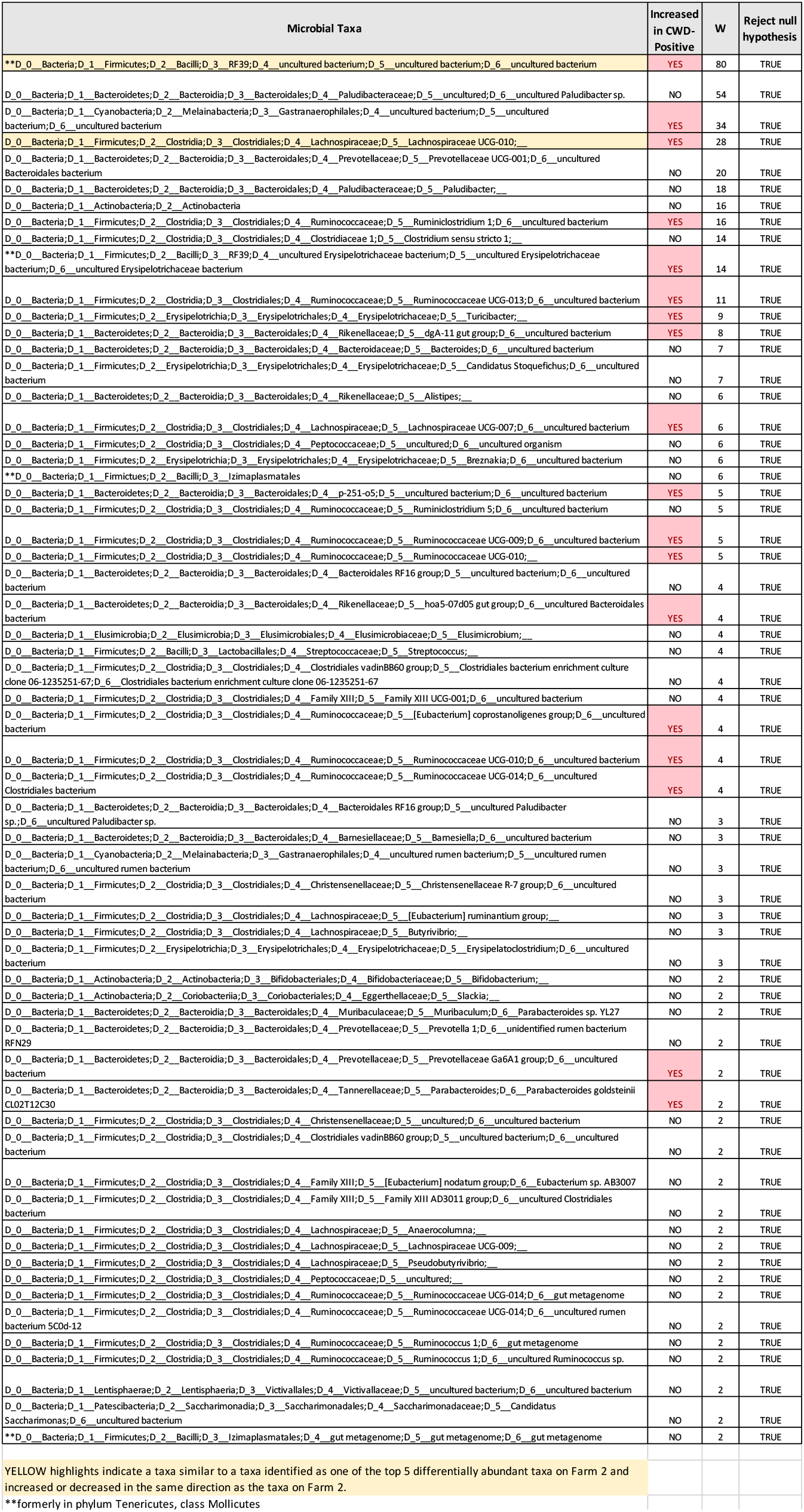
Differentially abundant taxa (ANCOM) by CWD Status on Farm 1 at the L7 (roughly species) level.

**Supplemental Table 4:**
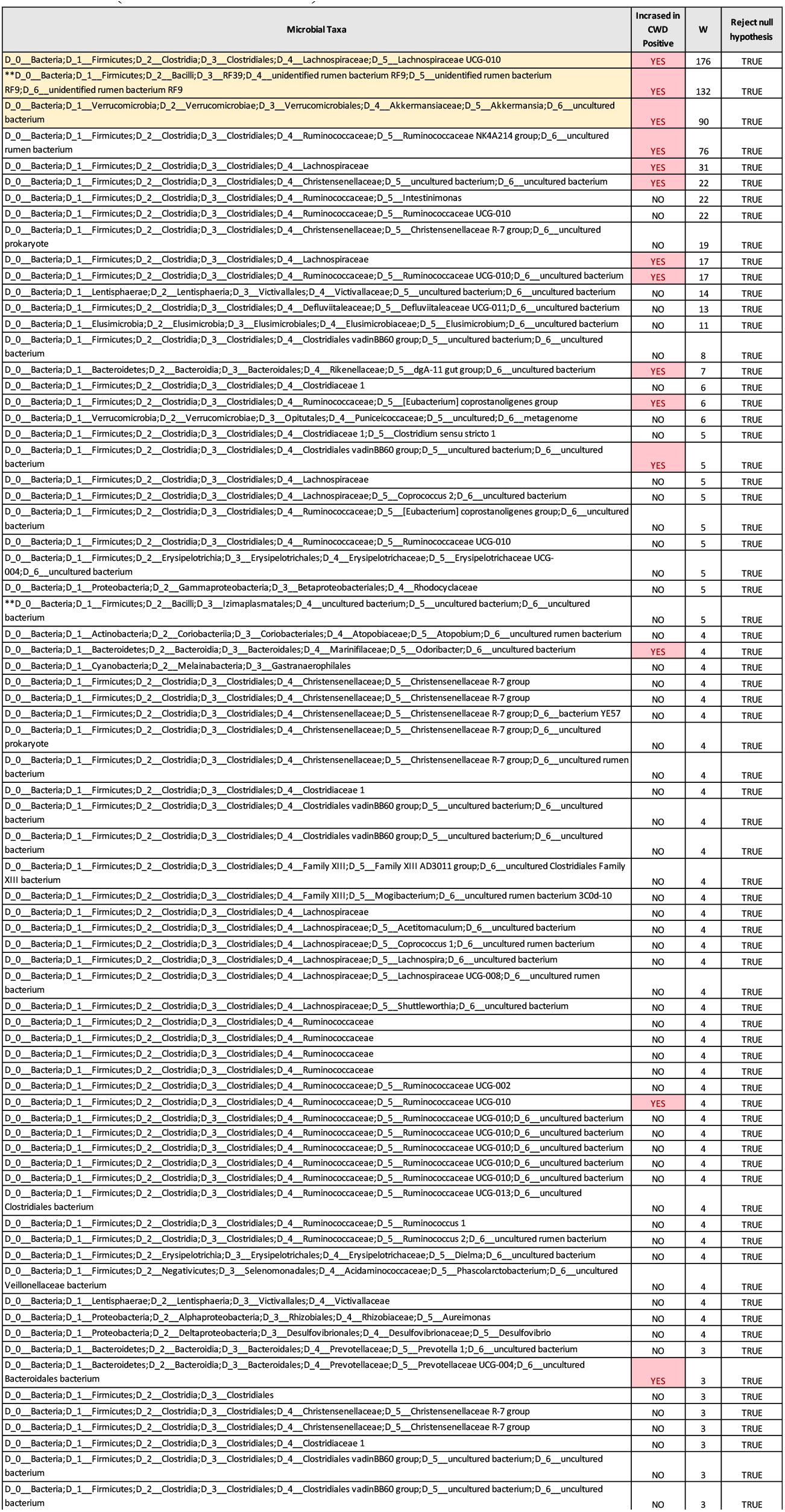
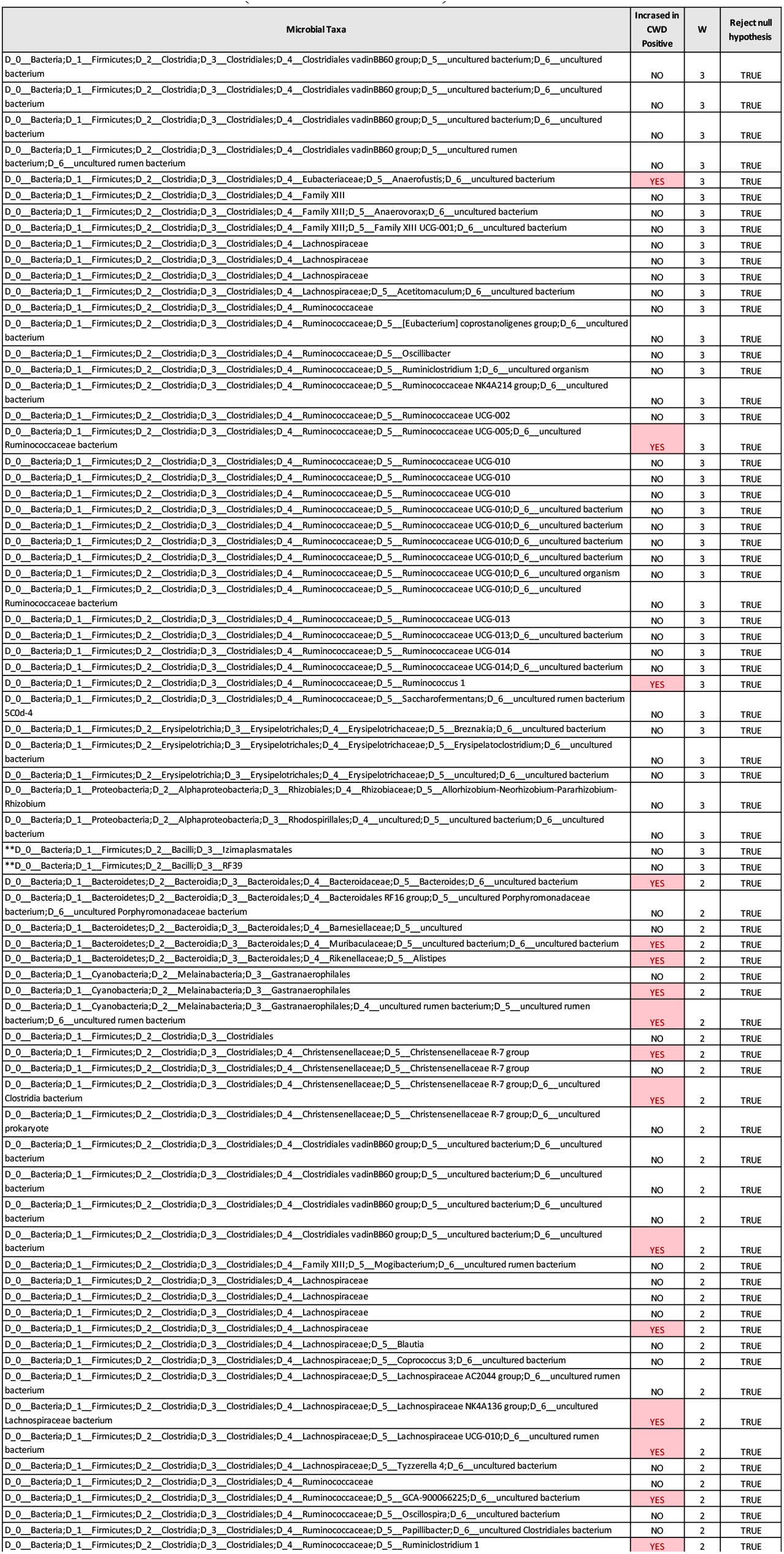
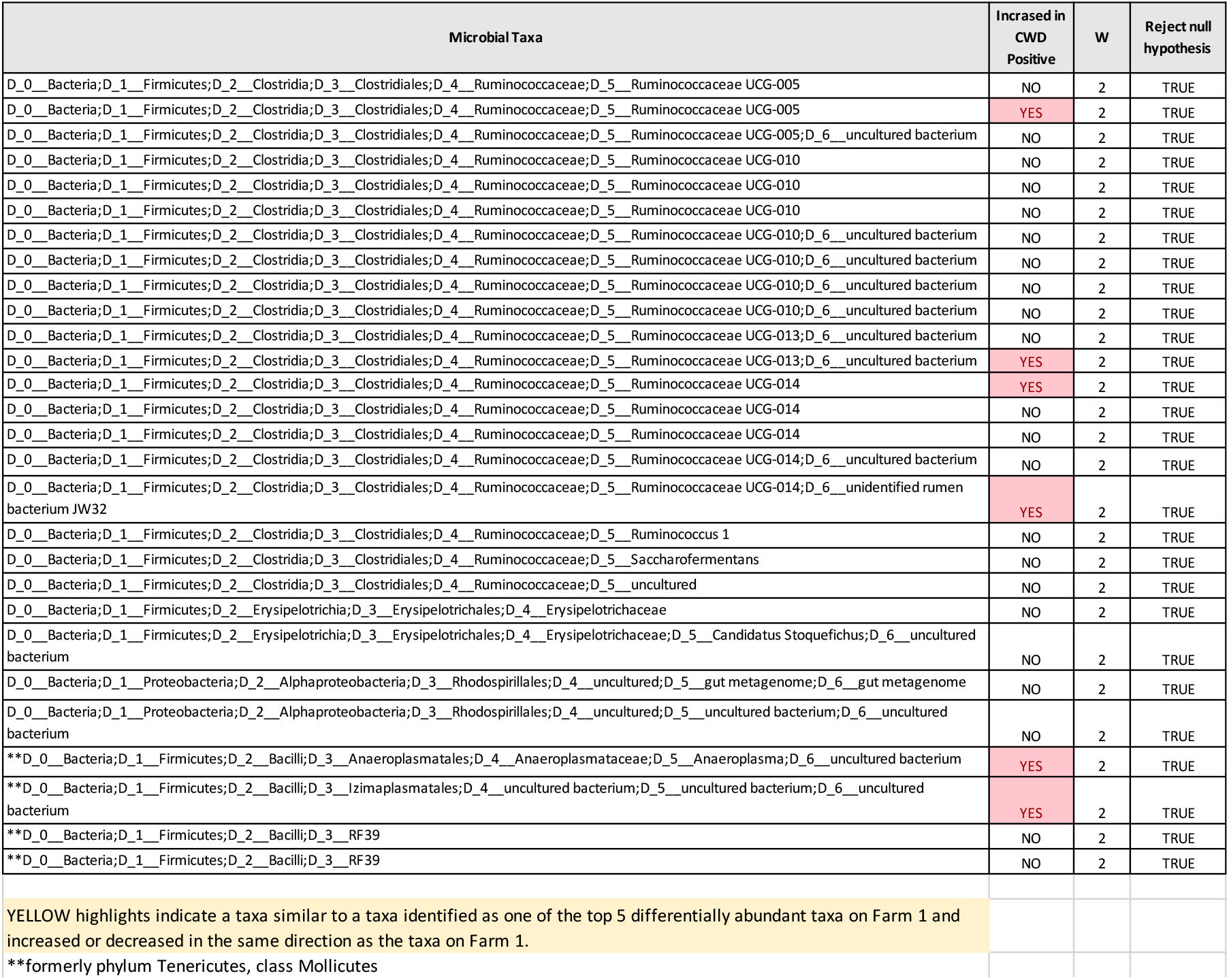
Differentially abundant taxa (ANCOM) by CWD Status on Farm 2 at the ASV (similar to strain) level.

## Notes

### Competing Interest Statement

The authors have declared no competing interest.

